# The ecological genetics of *Pseudomonas syringae* from kiwifruit leaves

**DOI:** 10.1101/235853

**Authors:** Christina Straub, Elena Colombi, Li Li, Hongwen Huang, Matthew D. Templeton, Honour C. McCann, Paul B. Rainey

## Abstract

Interactions between commensal microbes and invading pathogens are understudied, despite their likely effects on pathogen population structure and infection processes. We describe the population structure and genetic diversity of a broad range of co-occurring *Pseudomonas syringae* isolated from infected and uninfected kiwifruit during an outbreak of bleeding canker disease caused by *P. syringae* pv. *actinidiae* (*Psa*) in New Zealand. Overall population structure was clonal and affected by ecological factors including infection status and cultivar. Most isolates are members of a new clade in phylogroup 3 (PG3a), also present on kiwifruit leaves in China and Japan. Stability of the polymorphism between pathogenic *Psa* and commensal *P. syringae* PG3a isolated from the same leaf was tested using reciprocal invasion from rare assays *in vitro* and in planta. *P. syringae* G33C (PG3a) inhibited *Psa* NZ54, while the presence of *Psa* NZ54 enhanced the growth of *P. syringae* G33C. This effect could not be attributed to virulence activity encoded by the Type 3 secretion system of *Psa*. Together our data contribute toward the development of an ecological perspective on the genetic structure of pathogen populations.

**ORIGINALITY-SIGNIFICANT STATEMENT:** Bacterial pathogen populations are often studied with little consideration of co-occurring microbes and yet interactions between pathogens and commensals can affect both population structure and disease progression. A fine-scale sampling of commensals present on kiwifruit leaves during an outbreak of bleeding canker disease caused by *P. syringae* pv. *actinidiae* reveals a clonal population structure. A new clade of non-pathogenic *P. syringae* (PG3a) appears to be associated with kiwifruit on a global scale. The presence of PG3a on kiwifruit has significant effects on the outcome of infection by *P. syringae* pv. *actinidiae*. This emphasises the value of studying the effect of co-occurring bacteria on pathogen-plant interactions.

## INTRODUCTION

Kiwifruit (*Actinidia* spp.) cultivation is challenged by outbreaks of the bacterial pathogen *Pseudomonas syringae* pv. *actinidiae* (*Psa*) – the causative agent of bleeding canker disease. The latest outbreak was first reported in Italy in 2008 (Balestra *et al*., 2008) before spreading rapidly through most kiwifruit growing regions of the world (Abelleira *et al*., 2011; Everett *et al*., 2011; Koh *et al*., 2012; Zhao *et al*., 2013; Sawada, 2015), arriving in New Zealand in 2010 (Everett *et al*., 2011).

As a pathogen, *Psa* faces the challenge of colonising diverse environments before proliferating in the apoplast and vascular tissues. Colonisation of leaf surfaces prior to invasion is a key infection stage (Wilson and Lindow, 1994; Wilson *et al*., 1999; Monier and Lindow, 2003; Pfeilmeier *et al*., 2016). On the leaf surface *Psa* is likely to encounter and interact with a diverse range of plant-colonising bacteria (Hirano and Upper, 2000; Lindow and Brandl, 2003). Physical proximity increases the likelihood of competitive interactions affecting disease outcomes (Lindow and Brandl, 2003; Hibbing *et al*., 2010) and increases the probability of horizontal gene transfer (Sawada *et al*., 1999; Polz *et al*., 2013; Colombi *et al*., 2017).

The local context and the scale of sampling bacterial populations is particularly important, as it can have an impact on genetic structure (Istock *et al*., 1992; Souza *et al*., 1992; Haubold and Rainey, 1996; Spratt and Maiden, 1999). A study by Istock *et al*. (1992) made a particularly persuasive case by showing that *sampling Bacillus subtilis* at the local level (instead of pooled collections) contradicted the common view of its clonal population structure. Similar lessons regarding the scale of sampling have come from studying *P. syringae* populations with a range of structures reported depending upon whether or not environmental isolates are included (Sarkar and Guttman, 2004; Monteil *et al*., 2013).

Bacterial interactions are context-dependent, ranging from synergistic to antagonistic, and may have both local and global effects on the plant host (Stubbendieck *et al*., 2016). Antagonistic or competitive interactions between microbes may be direct or indirect, resulting in the inhibition of growth or even killing (Lindow, 1986; Völksch and May, 2001; Berlec, 2012; Hockett *et al*., 2015; Nakahara *et al*., 2016). Synergistic interactions occur when multiple types cooperate to cause disease (Singer, 2010; Lamichhane and Venturi, 2015). For example, *P. savastanoi* pv. *savastanoi*, causative agent of olive knot disease, interacts with non-pathogenic endophytes *Erwinia* sp. and sp. in cankers, enhancing the severity of disease (Marchi *et al*., 2006; Moretti *et al*., 2011; Buonaurio *et al*., 2015). Synergistic interactions can also be exploitative: bacteria lacking virulence factors can reap benefits from co-existing pathogenic isolates (Young, 1974; Hirano *et al*., 1999; Macho *et al*., 2007; Rufián *et al*., 2017).

*P. syringae* is a common member of the phyllosphere (defined as the aerial part of a plant (Vorholt, 2012)) and engages in both commensal and pathogenic interactions with plants (Hirano and Upper, 2000; Mohr *et al*., 2008). The diversity and population structure of *P. syringae* has been investigated using both multilocus sequence typing (MLST) and genome sequence analysis of pathogenic isolates collected from diseased plants (Sarkar and Guttman, 2004; Hwang *et al*.,2005; Baltrus *et al*., 2011; McCann *et al*., 2013, 2017; Fujikawa and Sawada, 2016; Nowell *et al*., 2016). Studies have also explored the structure of *P. syringae* populations from environmental reservoirs beyond standard host plants (Morris *et al*., 2008; Monteil *et al*., 2013, 2014, 2016). However, the genetic structure of specific pathovar populations from the phyllosphere of specific host plants have rarely been studied in the context of co-occurring *P. syringae* types.

Here we describe the population structure of the *P. syringae* species complex inhabiting the kiwifruit phyllosphere during an outbreak of bleeding canker disease in New Zealand. Using an MLST scheme, we reveal a largely clonal population structure, but show that genetic diversity is significantly affected by ecological factors such as infection status and cultivar. We identified members of four *P. syringae* phylogroups (PG1, PG2, PG3 and PG5) and recovered a new monophyletic clade within PG3 (PG3a) that is associated with kiwifruit in different kiwifruit-growing regions of the world. Investigations into the ecological interactions between a representative of this new clade and *Psa* show that PG3a restricts *Psa* proliferation, while *Psa* facilitates growth of PG3a.

## RESULTS

### Phyllosphere diversity of *Pseudomonas syringae*

Four housekeeping genes (*gapA, gyrB, gltA, rpoD*) were sequenced for each of 148 *P. syringae* isolated from two varieties of kiwifruit (‘Hayward’ and ‘Hort16A’) in both uninfected and *Psa*-infected orchards. Rarefaction analysis indicates saturation of the sampling effort (Figure S1). The infected ‘Hayward’ orchard displayed the highest α-diversity (D=0.904), while the uninfected ‘Hort16A’ orchard displayed the least α-diversity (D=0.737). There was low evenness (ED) among all sampling sites (0.136 to 0.290). Similarly, the four different sampling sites shared few species (Sørensen’s index of dissimilarity = 0.847).

### Multilocus sequence typing

45 unique sequence types (ST) were discovered among the 148 sequenced strains. All STs were novel, except for ST904 (*Psa*), and not described in the Plant Associated and Environmental Microbes Database (PAMDB). For a more global analysis, *P. syringae* allelic profiles were sourced from PAMDB and NCBI (accessions listed in Table S2), along with *P. syringae* isolated from kiwifruit in Japan (NCBI), NZ and the US (Visnovsky *et al*. 2016).

Infected orchards (both ‘Hayward’ and ‘Hort16A’) harboured the highest number of unique STs, sharing only three STs between them (Figure S2). No STs were present in all four orchards, but two STs were found in three orchards (ST1 and ST3). From the perspective of clonal complexes (CC), the predominant ST (predicted founder) was present along with several SLVs (single locus variants). Two clonal complexes (CC) (21 strains), 5 doubletons (32 strains) and 28 singletons (95 strains) were identified (Figure S3). CC1 and CC2 are comprised of 11 and 10 strains, respectively. ST904 (*Psa*, 15/148) and ST1 (PG3, 24/148) made up 25% of the sample (Figure S3). Strikingly, ST3 (PG3a) was isolated from three out of four orchards and was also sampled from uninfected gold (A. *chinensis* var. *chinensis*) and green (*A.chinensis* var. *deliciosa*) kiwifruit in NZ (2010) (Visnovsky *et al*., 2016) and Japan (2015), respectively (Figure 1). ST16 was also recovered from uninfected kiwifruit leaves in NZ in both 1991 and 2013. Other Japanese kiwifruit STs group closely with P.syringae originating from kiwifruit in NZ.

**Figure 1.**
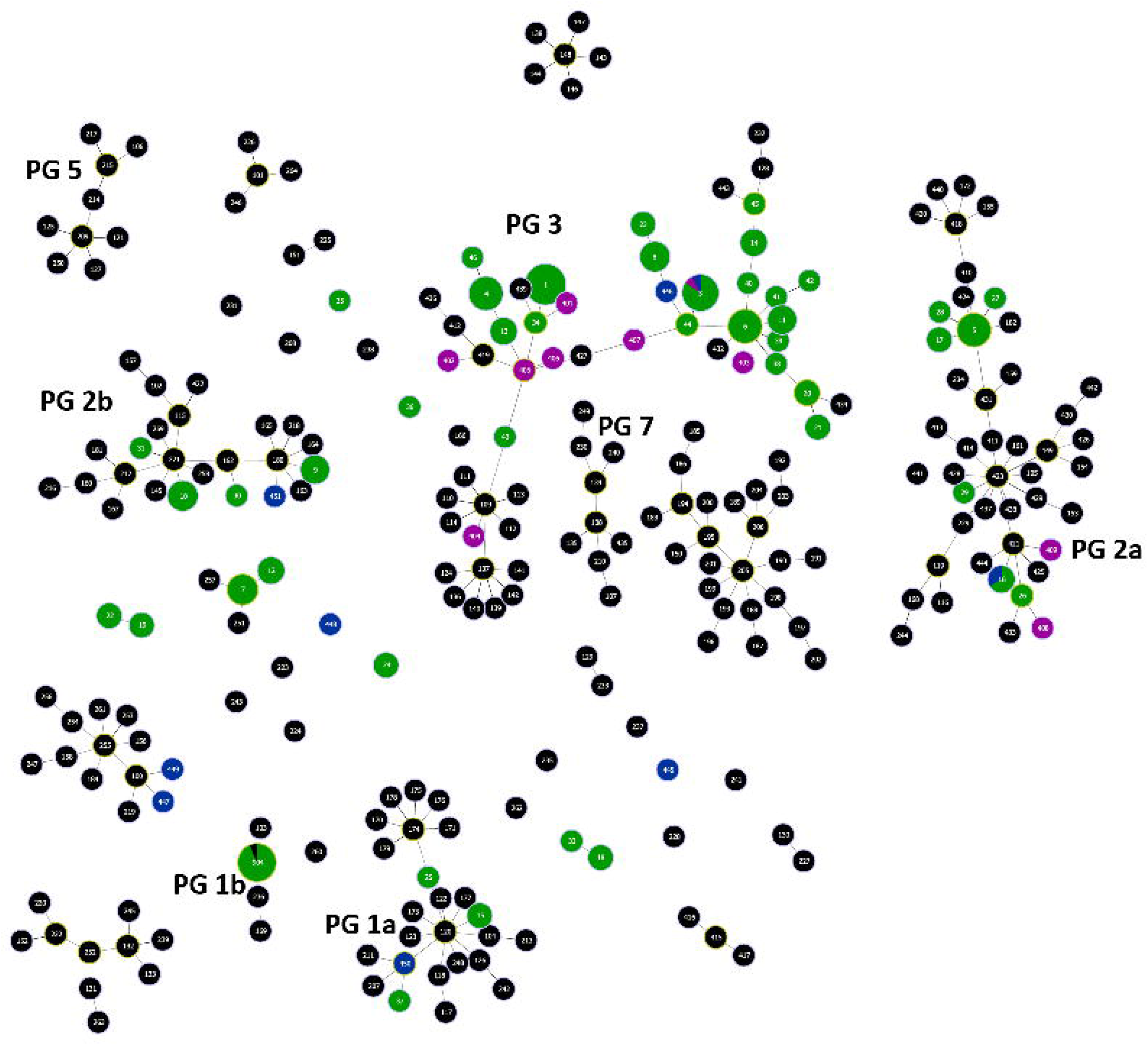
Global Minimum Spanning Tree (MST). Displaying the relationships between STs at the triple-locus-variant level, illustrated using PHYLOViZ (Francisco *et al.*, 2012). The size of the circle correlates with the frequency of the ST. Color-coded accorded to origin: green = this study, black = PAMDB, blue = Visnovsky *et al.* (2016), purple = Japan.

### Sequence diversity

The total concatenated alignment length was 2010 bp with no insertions or deletions detected for either of the four loci. The number of alleles ranged from 25 (*gapA*) to 35 (*rpoD*) (Table 1). There were a total of 412 polymorphic sites, ranging from 80 (16.81%, *gapA*) to 145 (28.6%, *gyrB*). The nucleotide diversity index π and Watterson’s θ were highly consistent among loci, varying from 0.040 to 0.055 and 0.024 to 0.041 respectively. The average GC content of 57.99% is similar to that found in other *P. syringae* studies (59-61%). The pairwise genetic difference within phylogroups (PGs) was not greater than 2.7%, whereas among PGs the variability ranged from 6-11% (Table 2), consistent with previous accounts of genetic variability for *P. syringae* (Sarkar and Guttman, 2004; Morris *et al*., 2010; Berge *et al*., 2014).

**Table 1:** Nucleotide and amino acid diversity. L = length in bp, AA = amino acid, GC = average GC content in %, N_A_ = number of alleles, P = number of polymorphic sites, *d*_*N*_/*d*_*S*_ ration, mut = mutations, π = nucleotide diversity indices, θ = Watterson’s theta.

| Locus | L<br>(bp) | AA<br>length | GC | $N_A$ | P* | $d_N/d_S$ | mut | $\pi$ | $\theta$ |
| --- | --- | --- | --- | --- | --- | --- | --- | --- | --- |
| *gapA* | 476 | 158 | 60.81 | 25 | 80 (16.81) | 3.365 | 97 | 0.055 | 0.024 |
| *gyrB* | 507 | 169 | 53.15 | 28 | 145 (28.60) | 0.018 | 184 | 0.054 | 0.041 |
| *gltA* | 529 | 176 | 58.45 | 27 | 88 (16.64) | 0.011 | 102 | 0.040 | 0.025 |
| *rpoD* | 495 | 166 | 59.56 | 35 | 99 (20) | 2.022 | 118 | 0.042 | 0.030 |
| Mean | 502 | 167 | 57.99 | 29 | 105 (20.51) | 1.354 | 125 | 0.048 | 0.030 |

**Table 2:**
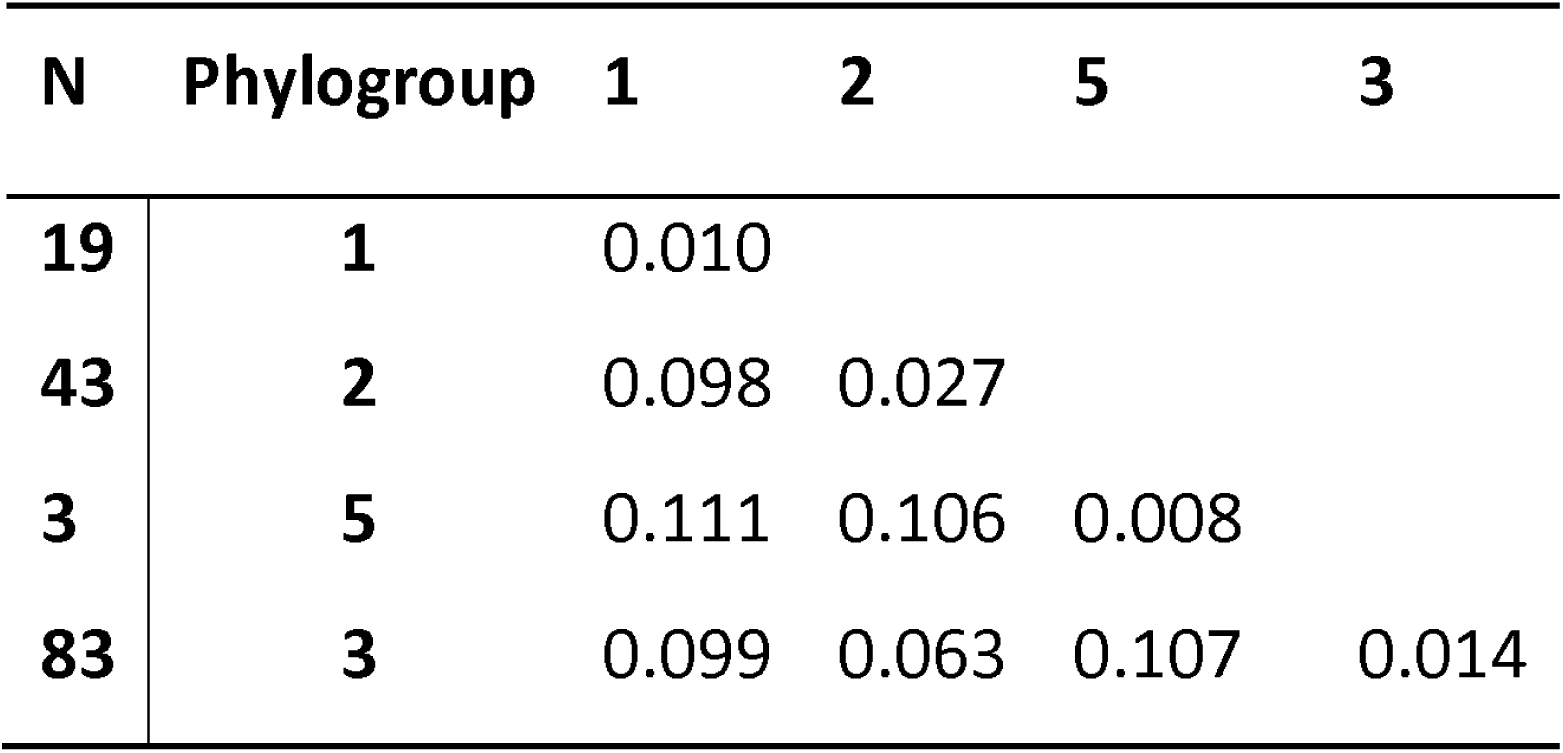
Average pairwise genetic diversity between and among phylogroups. Analyses were conducted on the concatenated alignment (2006 bp, gaps removed) using the Maximum Composite Likelihood model with a gamma distribution of 1. N= number of strains.

| N | Phylogroup | 1 | 2 | 5 | 3 |
| --- | --- | --- | --- | --- | --- |
| 19 | 1 | 0.010 |  |  |  |
| 43 | 2 | 0.098 | 0.027 |  |  |
| 3 | 5 | 0.111 | 0.106 | 0.008 |  |
| 83 | 3 | 0.099 | 0.063 | 0.107 | 0.014 |

### Genetic diversity varies by host cultivar and infection status

To test whether genetic diversity was influenced by the presence of ecological structure, multivariate analyses (PERMANOVA) were performed. A highly significant difference in genetic diversity was observed among sampled orchards (Pseudo-F = 5.99, *P <* 0.0001); with pairwise Permanova tests revealing that the uninfected green orchard differed significantly from every other orchard (*P*<0.003). The cultivar (‘Hayward’ vs. ‘Hort16A’) (Pseudo-F = 5.62, *P*<0.001) and infection status of an orchard (Pseudo-F = 11.72, *P* <0.001) also had a significant effect on genetic diversity, whereas no temporal effect was found (Pseudo-F = 1.10, *P* >0.34). When testing the nested effect of all three factors, only the infection status (Pseudo-F = 6.42, *P* <0.01) had a significant impact on genetic diversity (Table S3).

### Recombination among *P. syringae*

Intragenic recombination rates (ρ) ranged from 0.012 (*rpoD*) to 0.038 (*gyrB*) and 0.006 for the concatenated dataset. The ratio ε (recombination rate/mutation rate) ranged from 0.187 (concatenated) to 0.931 (*gyrB*) suggesting that any single nucleotide polymorphism is up to five times more likely to have arisen from a mutation than recombination (Table 3).

**Table 3:** LDhat recombination analysis. Showing the length of the alignment in bp, N = number of sequences, mutation rate θ (=2Neμ) per site, recombination rate ρ (=2Ner) per site, ratio ε = ρ/ θ and Tajima’s D.

| Gene | Length (bp) | N | Segregating sites | $\theta$ | $\rho$ | $\epsilon = \rho / \theta$ | Tajima's D |
| --- | --- | --- | --- | --- | --- | --- | --- |
| **All *P. syringae*:** |  |  |  |  |  |  |  |
| concatenated | 2010 | 148 | 335 | 0.030 | 0.006 | 0.187 | 0.513 |
| *gapA* | 476 | 148 | 63 | 0.024 | 0.021 | 0.902 | 2.204 |
| *gyrB* | 507 | 148 | 116 | 0.041 | 0.038 | 0.931 | -0.204 |
| *gltA* | 529 | 148 | 74 | 0.025 | 0.012 | 0.461 | 0.678 |
| *rpoD* | 498 | 148 | 82 | 0.030 | 0.012 | 0.416 | 0.03 |
| **Phylogroup 1** |  |  |  |  |  |  |  |
| concat | 2010 | 19 | 66 | 0.009 | 0.000 | 0.000 | 0.207 |
| *gapA* | 476 | 19 | 8 | 0.005 | 0.000 | 0.000 | 0.407 |
| *gyrB* | 507 | 19 | 20 | 0.011 | 0.000 | 0.000 | 0.611 |
| *gltA* | 529 | 19 | 19 | 0.010 | 0.000 | 0.000 | 0.026 |
| *rpoD* | 498 | 19 | 19 | 0.011 | 0.000 | 0.000 | -0.188 |
| **Phylogroup 2** |  |  |  |  |  |  |  |
| concat | 2010 | 43 | 147 | 0.017 | 0.002 | 0.120 | 1.754 |
| *gapA* | 476 | 43 | 29 | 0.014 | 0.006 | 0.457 | 2.307 |
| *gyrB* | 507 | 43 | 63 | 0.029 | 0.000 | 0.000 | 2.131 |
| *gltA* | 529 | 43 | 27 | 0.012 | 0.008 | 0.654 | 0.993 |
| *rpoD* | 498 | 43 | 28 | 0.013 | 0.023 | 1.734 | 0.621 |
| **Phylogroup 3** |  |  |  |  |  |  |  |
| concat | 2010 | 83 | 163 | 0.016 | 0.015 | 0.937 | -0.746 |
| *gapA* | 476 | 83 | 34 | 0.014 | 0.013 | 0.898 | 2.096 |
| *gyrB* | 507 | 83 | 96 | 0.038 | 0.024 | 0.636 | -2.434 |
| *gltA* | 529 | 83 | 13 | 0.005 | 0.006 | 1.175 | 1.991 |
| *rpoD* | 498 | 83 | 20 | 0.008 | 0.033 | 4.073 | 0.3 |
| **Phylogroup 5** |  |  |  |  |  |  |  |
| concat | 2010 | 3 | 23 | 0.008 | 0.000 | 0.000 | - |
| *gapA* | 476 | 3 | - | - | - | - | - |
| *gyrB* | 507 | 3 | 13 | 0.017 | 0.000 | 0.000 | - |
| *gltA* | 529 | 3 | 3 | 0.004 | 0.000 | 0.000 | - |
| *rpoD* | 498 | 3 | 7 | 0.009 | 0.000 | 0.000 | - |

Clustering sequences by PG revealed no evidence of recombination within PG1 and PG5 (ρ=0), however for these phylogroups the sample size was low. There was evidence of recombination in PG3, more specifically for *gltA* (ε=1.18) and *rpoD* (ε=4.07), whereas in PG2 recombination was evident in *rpoD* (ε=1.734) alone. Recombination was neither affected by the host cultivar or infection status (Table S4). The analysis was repeated with the inclusion of non-redundant global strains. Overall, intergenic recombination rates were low among PGs, ranging from 0.005 to 0.012 for the concatenated dataset (Table S5). In order to pinpoint any effects of recombination on phylogenetic reconstruction, single gene trees were constructed and compared. Tree topologies were significantly different from each other and from the concatenated dataset (SH test, *P*<0.05) (Figure S4).

### A kiwifruit-associated clade of *P. syringae*

Maximum likelihood trees built using the concatenated alignment of unique STs revealed that nearly all NZ *P. syringae* kiwifruit isolates fell within four PGs: PG1 (13%), PG2 (29%) and PG3 (56%), with only a few isolates falling into PG5 (2%, Figure 2). Surprisingly, within PG3 all NZ kiwifruit-associated isolates grouped within a new clade of PG3, hereafter referred to as PG3a. The uninfected orchards showed a higher number of PG3a isolates compared to infected orchards, although no influence of infection status was reflected in the number of unique sequence types (Table S1). We also found that two strains isolated from kiwifruit in NZ in 2010 and 2011 (Visnovsky *et al*., 2016) belong to this subclade.

**Figure 2.**
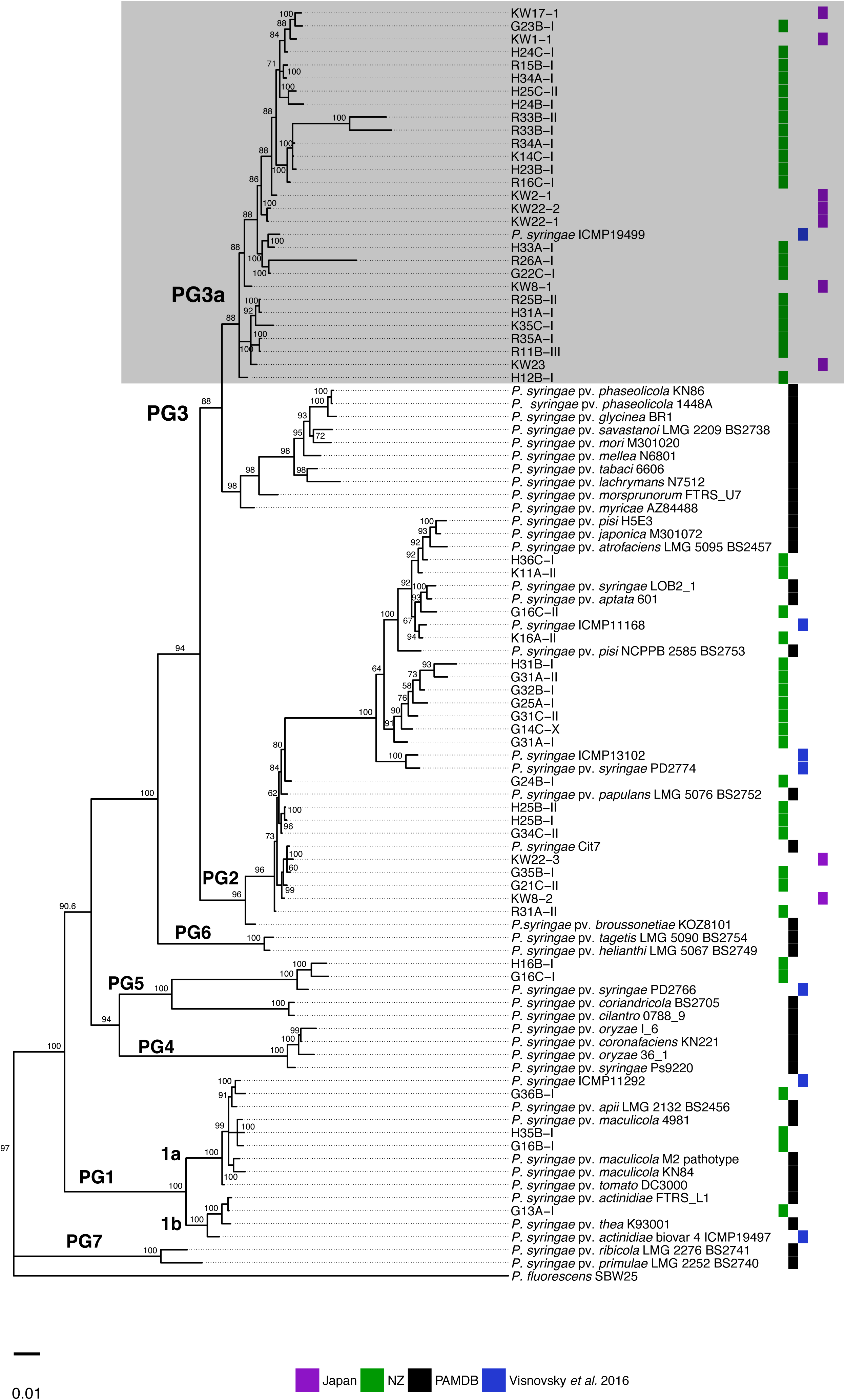
Bayesian tree based on the concatenated alignment (2010 bp) of four housekeeping genes: *gapA, gyrB, gltA* and *rpoD*. The Bayesian tree was reconstructed using MrBayes based on the Tamura-Nei + G + I model using 30,000,000 MCMC. Single representative sequences for each ST were used to improve readability (corresponding STs and frequency of each ST listed in Table S2). Values indicated at nodes are Bayesian posterior probabilities. The corresponding phylogroups (PG) are indicated, eg. PG1 = phylogroup 1 with clades 1a and 1b. Origin of isolates is illustrated in colour coded boxes, green = this study, black = PAMDB, blue = Visnovsky *et al.* 2016, purple = Japan.

This discovery led us to question whether the PG3a subclade might be prevalent on kiwifruit vines in other countries. To this end we interrogated an unpublished set of *gltA* sequences from *P. syringae* strains collected from kiwifruit leaves in China (sampling as described in McCann *et al*. (2017), GenBank accession numbers: MG674624 – MG674645). A phylogenetic tree based on *gltA* for NZ, Japanese and Chinese kiwifruit isolates revealed that isolates obtained from Chinese kiwifruit also clustered within PG3a (Figure 3). Interestingly, included in this group is isolate 47L9, which was collected from tea leaves (Camellia sp.) growing in a former kiwifruit orchard in China. No other *P. syringae* strain from the PAMDB database grouped with PG3a, suggesting PG3a is persistently associated with kiwifruit on a global scale.

**Figure 3.**
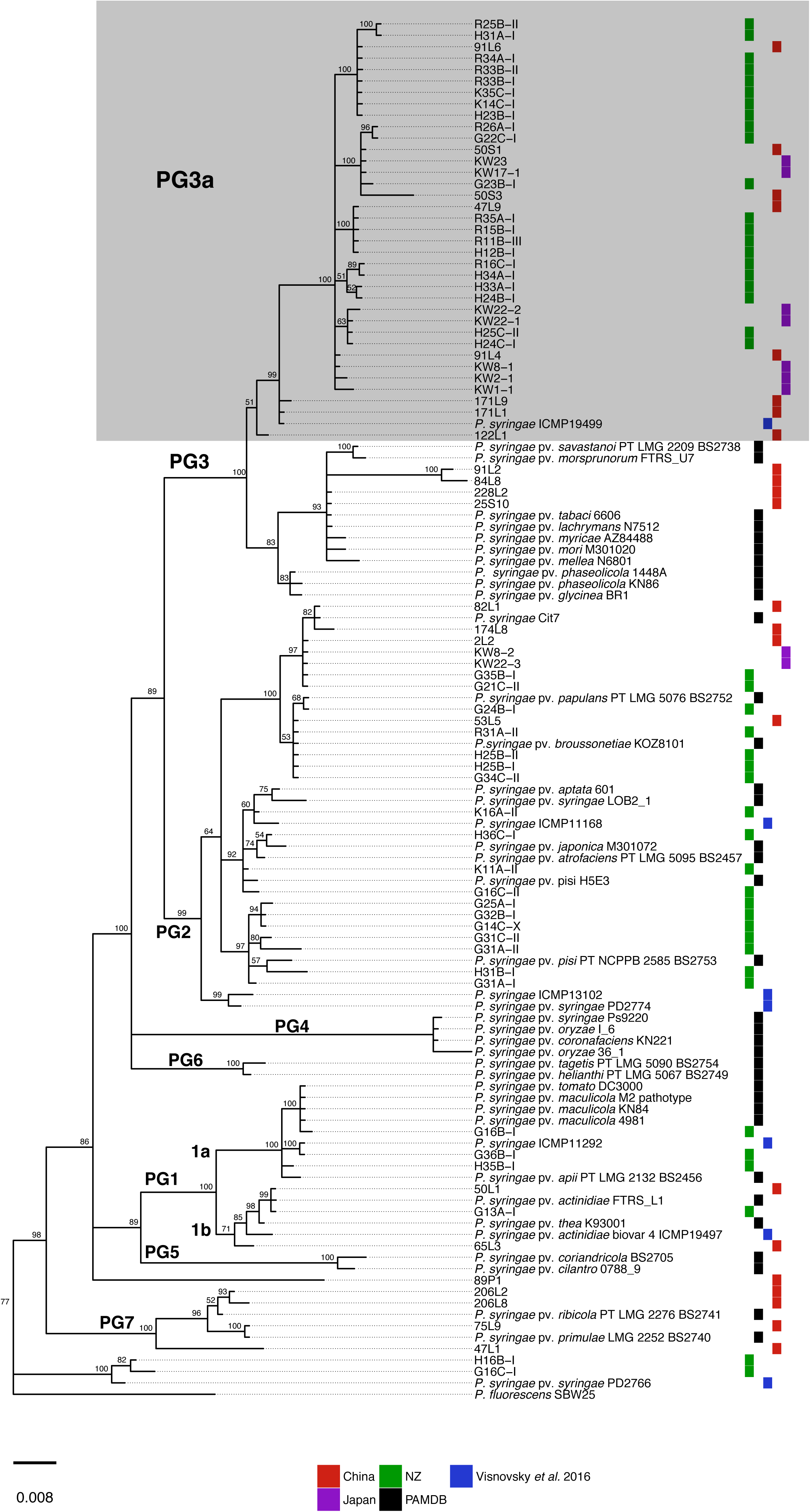
Global Bayesian tree reconstructed from *gltA* sequences highlighting the particularity of PG3a, which includes kiwifruit isolates from NZ, China, Japan, the US and France. The tree was built on a 529 bp alignment using MrBayes (TN+G+I model; 1,700,000 MCMC), using *Pseudomonas fluorescens* SBW25 as outgroup. Values indicated at nodes are Bayesian posterior probabilities. The source of each isolate is highlighted in colour-coded boxes, green = this study, red = China, black = PAMDB, purple = Japan, blue = US, NZ and France.

### Ecological interactions between PG3a and PG1

To assess whether kiwifruit-associated PG3a strains are stably maintained with *Psa* (PG1), co-inoculation experiments were performed *in vitro* and in planta. Two isolates sampled from the same leaf were chosen for these experiments: *P. syringae* G33C (ST1, PG3a) and *Psa* NZ54 (ST904, PG1).

### In vitro *dynamics*

*Psa* NZ54 and *P. syringae* G33C showed similar growth dynamics when grown individually *in vitro* (Figure 4). However, when co-inoculated in liquid King’s B (KB) media at an equal starting ratio, *Psa* NZ54 growth was significantly reduced (up to 100-fold) at 24 h (*P* <0.001, paired *t*-tests). This effect was amplified in shaken liquid minimal M9 medium, which better approximates nutrient poor conditions in the leaf compared to KB medium (Hernández-Morales *et al*., 2009): *Psa* NZ54 population density collapsed by 20 h in shaken M9 media (relative fitness −12.1 ±0.09, Figure 4).

**Figure 4.**
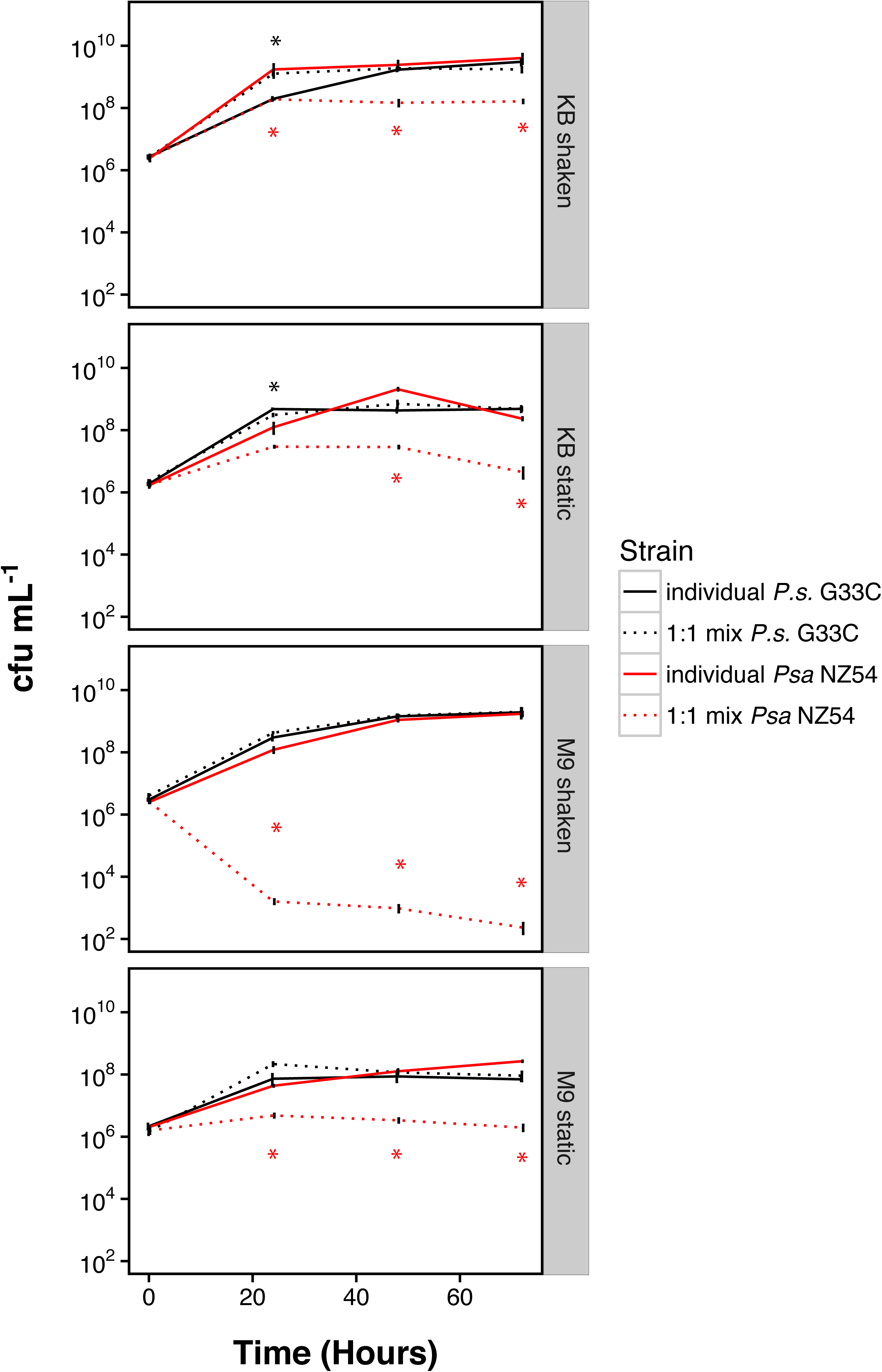
Individual growth dynamics of *Psa* NZ54 and *P. syringae* G33C compared with co-inoculation (1:1 ratio) *in vitro*. Competition experiments were performed in a 1:1 ratio (founding ratio 5×10^6^ cfu ml^−1^ each), with individual inoculations as reference. Solid lines represent individual growth and dashed lines represent growth in competition. The presented mean and standard error were calculated from three replicates. Asterisk indicate significance between individual and co-cultured growth at the 5% level (paired *t-*test).

In order to establish whether the instability of the interaction was influenced by the ratio of founder cells, we investigated whether *P. syringae* G33C could invade from rare initial frequency. A strain is able to invade from rare when it increases relative to its founding density. *P. syringae* G33C successfully invaded from rare after only 24 h in both rich KB and minimal M9 media (10:1 *Psa* NZ54: *P. syringae* G33C) and reached a similar population size as when cultured on its own in M9 (Figure 5A). Conversely, *Psa* NZ54 also invaded *P. syringae* G33C from rare (1:10 *Psa* NZ54: *P. syringae* G33C), though it established a 100-fold reduced population size of 10^5^ - 10^6^ cfu ml^−1^ compared to growth alone. The population collapse of *Psa* NZ54 in shaken M9, as observed for the 1:1 competition experiments, was once again observed (Figure 5B). The striking suppression of *Psa* NZ54 by *P. syringae* G33C was unambiguously repeated across three experiments. *P. syringae* G33C outcompetes *Psa* NZ54, though both isolates can invade from rare *in vitro* (with the exception of *Psa* NZ54 in shaken M9), which suggests that in a controlled environment the polymorphism is stable.

**Figure 5.**
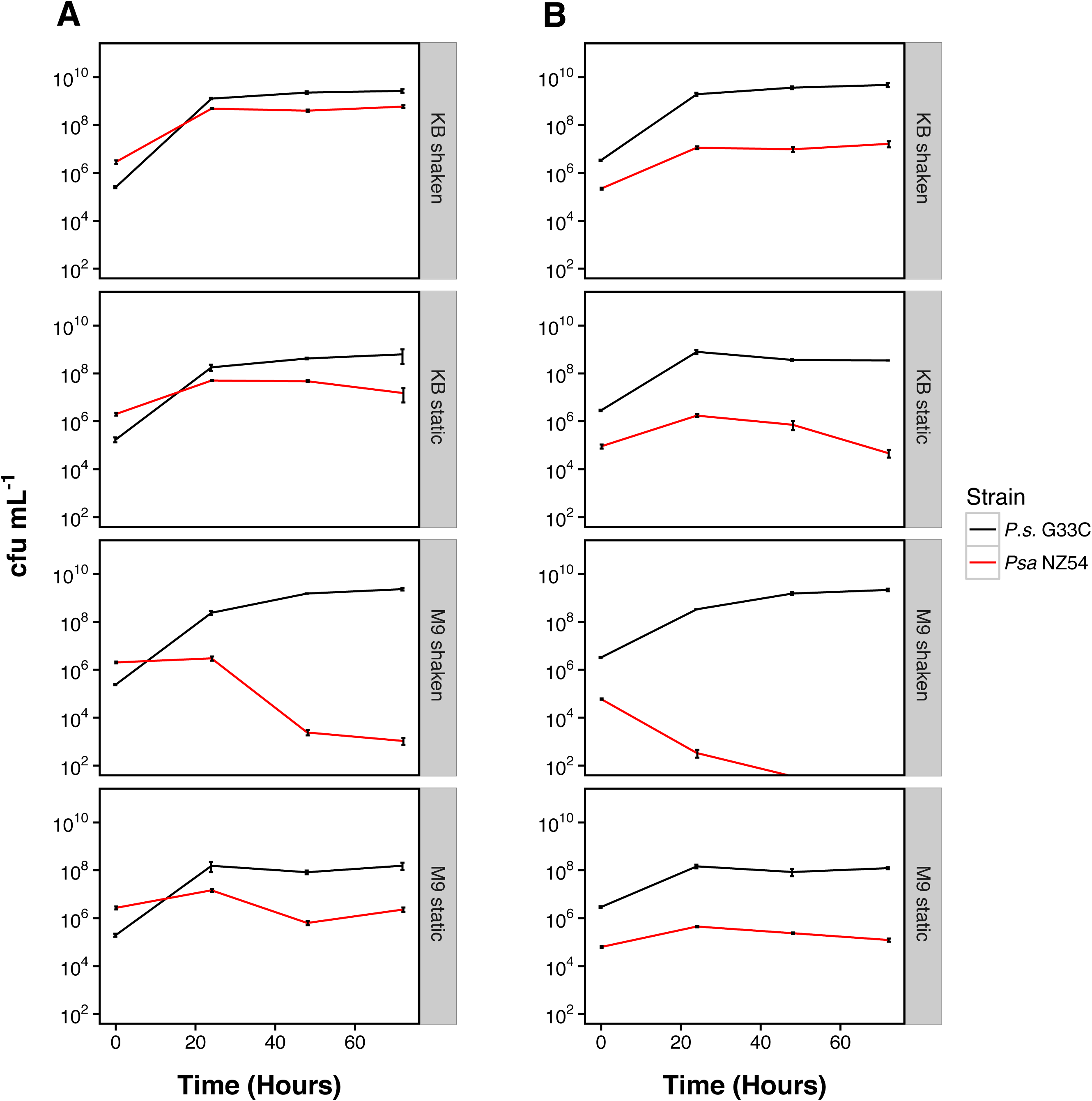
*In vitro* growth curves from invasion from rare experiments for *Psa* NZ54: *P. syringae* G33C and vice versa. Vials were inoculated with a (A) 10:1 ratio and (B) 1:10 ratio for *Psa* NZ54: *P. syringae* G33C. The presented mean and standard error were calculated from three replicates. Parameters of relative fitness of *Psa* NZ54 relative to *P. syringae* G33C calculated as ln difference *Psa* NZ54 – *P. syringae* G33C using the Malthusian parameters at 24hrs were −2.1* ±0.02 (KB shaken), −1.9*±0.05 (KB static), −12.1*±0.09 (M9 shaken) and −3.9*±0.00 (M9 static). Asterisks indicate significance at the 1% level (Students *t*-test).

### In planta *dynamics*

*In planta* experiments were performed on two gold cultivars, ‘Hort16A’ and ‘SunGold’, to determine whether *Psa* NZ54 and *P. syringae* G33C also form a stable polymorphism on kiwifruit leaves. *P. syringae* G33C established an epiphytic and endophytic population size in both cultivars (Figure 6) and did not produce any visible symptoms in ‘Hort16A’ and ‘SunGold’ (Figure S5, Figure S6). *Psa* NZ54 attained a population size at least 10,000-fold greater than *P. syringae* G33C in both hosts. However, endophytic and epiphytic growth were reduced 10-fold in ‘SunGold’ compared to ‘Hort16A’ (*P* <0.05, Mann-Whitney U test). Plants inoculated with *Psa* NZ54 developed the first leaf spots at 4 dpi and exhibited severe symptoms at 7 dpi in the more susceptible ‘Hort16A’, whereas in ‘SunGold’ leaves displayed only minor symptoms at 7 dpi.

**Figure 6.**
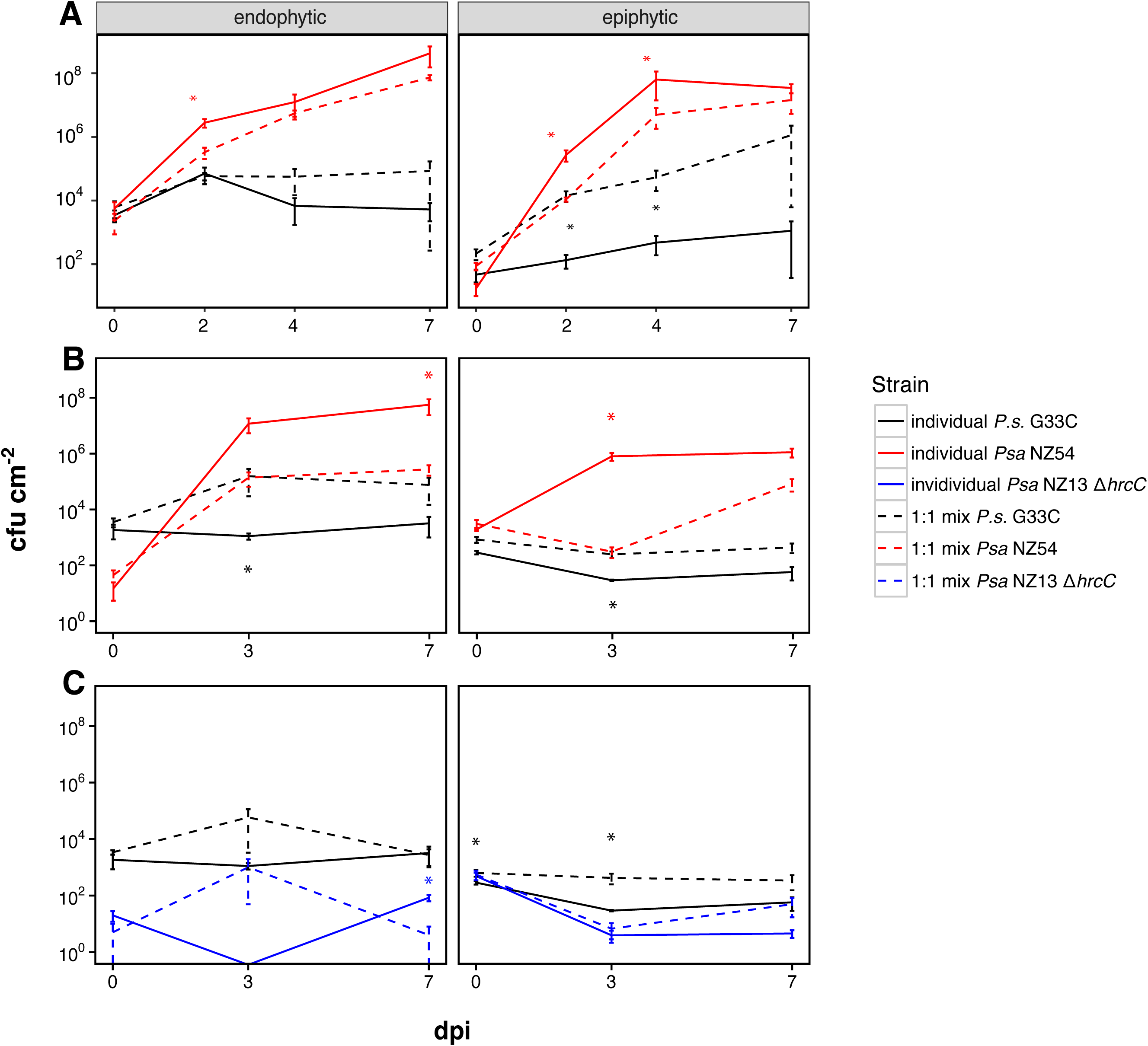
1:1 competition growth assays of *Psa* NZ54 vs. *P. syringae* G33C *in planta*. ‘Hort16A’ plantlets (A) and ‘SunGold’ plantlets (B) were inoculated with a 1:1 mix of *P. syringae* G33C: *Psa* NZ54 (founding density 8×10^7^ cfu ml^−1^). (C) ‘SunGold’ plants were inoculated with 1:1 mix of *P. syringae* G33C: *Psa* NZ13 Δ*hrcC* (founding density 8×10^7^ cfu ml^−1^). Solid lines represent individual growth and dashed lines represent growth in competition. The presented mean and standard error were calculated from the mean of four (‘Hort16A’) and five (‘SunGold’) individual measurements. Asterisk indicate significance between individual and co-cultured growth at the 5% level (paired *t-*test).

In 1:1 competition experiments non-pathogenic *P. syringae* G33C maintained a stable population size. The presence of *Psa* NZ54 had a highly significant positive effect on the growth of *P. syringae* G33C in both plant hosts (Figure 6A&B). *P. syringae* G33C established up to 1000-fold higher epiphytic population densities in ‘Hort16A’ (*P* <0.01, paired *t*-tests) and 10-fold higher epiphytic and endophytic population densities in ‘SunGold&’ plants (*P* <0.05, paired *t*-test) compared to its individual growth. Co-inoculated *Psa* NZ54 exhibited a significant reduction (*P* <0.05, paired *t*-test) in epiphytic and endophytic growth on ‘Hort16A’ in the presence of *P. syringae* G33C, but only in the early stages of the experiment. On ‘SunGold’ the diminished growth of *Psa* NZ54 was more pronounced, with a 100-fold decrease for the endophytic population at 7 dpi (*P* <0.05, paired *t*-test, Figure 6B). Co-inoculated ‘Hort16A’ plants exhibited a notable delay in symptom onset compared to singly inoculated plants (Figure S5), whereas there was no difference for ‘SunGold’ (Figure S6). The increased fitness of *Psa* NZ54 relative to *P. syringae* G33C in ‘Hort16A’ competition experiments was reflected in the relative fitness parameters (0.7± 0.1* for epiphytic and 4.9± 0.8* for endophytic, * *P* <0.05, *t*-test), whereas in ‘SunGold’ plants *P. syringae* G33C performed better in the epiphytic environment (−1.6 ± 0.2; 4.7 ± 0.1* for endophytic growth).

To assess whether the heightened growth of *P. syringae* G33C in the presence of *Psa* NZ54 was due to the virulence activity of the pathogen elicited by the Type 3 Secretion System (T3SS), the competition experiment was performed using a T3SS deficient mutant (*Psa* NZ13 Δ*hrcC*). Epiphytic growth of *P. syringae* G33C on ‘SunGold’ remained elevated when co-inoculated with *Psa* NZ13 Δ*hrcC*, indicating the virulence activity encoded by the T3SS was not responsible for the advantage conferred to the non-pathogenic strain (*P* <0.05, paired *t*-test, Figure 6C).

A rarity threshold for *P. syringae* G33C determines the ability to establish a stable population in planta. Upon co-inoculation in a 100:1 (*Psa* NZ54: *P. syringae* G33C) ratio on ‘Hort16A’, *P. syringae* G33C was able to invade from rare over the first 4 days, but was then excluded by *Psa* NZ54 (Figure 7B). An initial increase in growth of *P. syringae* G33C from 0 dpi to 4 dpi was followed by a population collapse at 7 dpi with no endophytic growth detected and minimal epiphytic growth in environments dominated by *Psa* NZ54. Conversely, *Psa* NZ54 grew to the same population size in the presence of *P. syringae* G33C, as when inoculated individually (*P* >0.1, paired *t*-tests).

**Figure 7.**
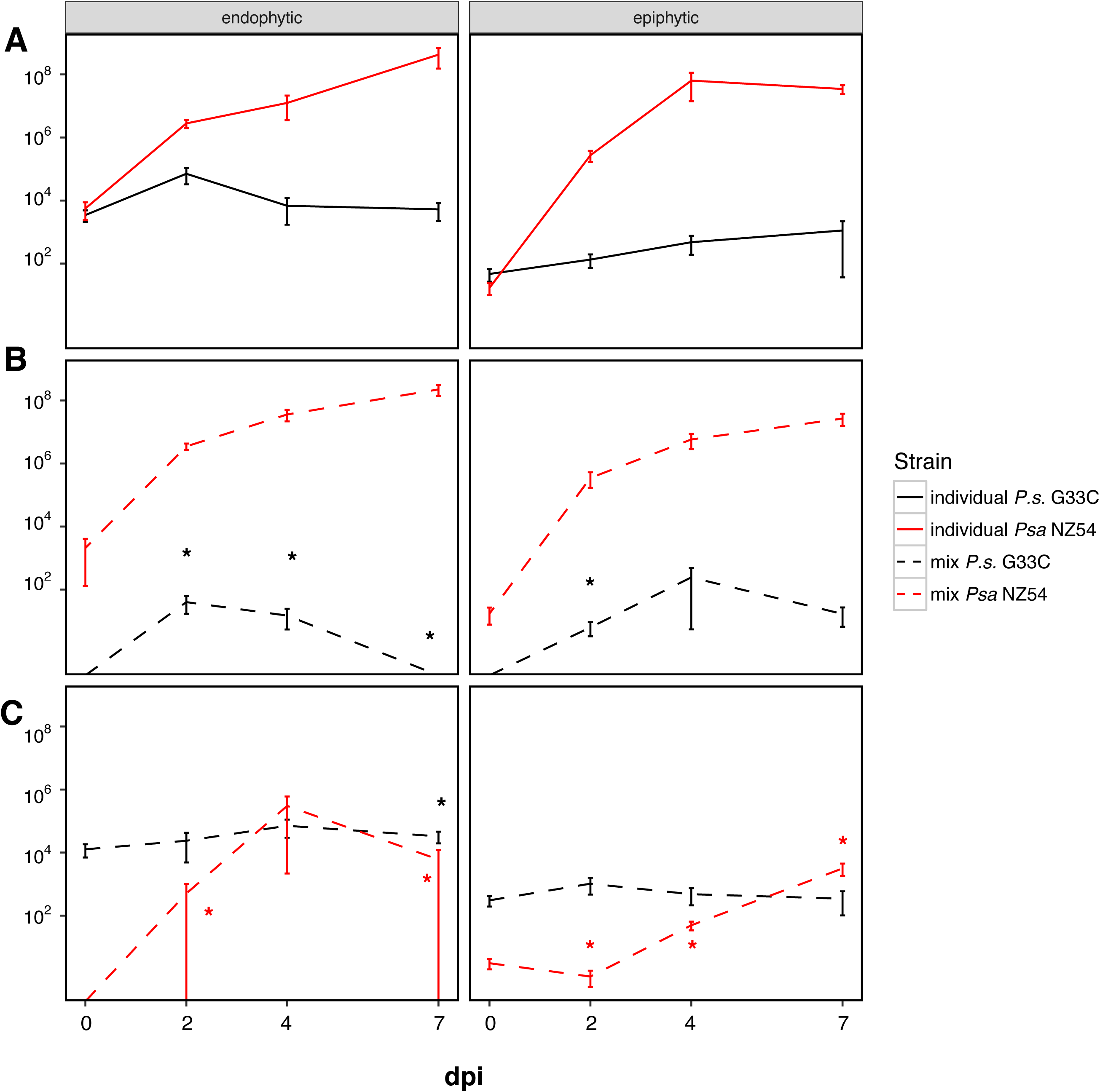
Invasion from rare experiments for *Psa* NZ54: *P. syringae* G33C *in planta*. ‘Hort16A’ plantlets were inoculated with different ratios of strains *Psa* NZ54: *P. syringae* G33C. A) Individual growth. B) Invasion from rare 100:1 and C) invasion from rare 1:100. Solid lines represent individual growth and dashed lines represent growth in competition. The presented mean and standard error were calculated from the mean of four individual measurements. Asterisks indicate significance between individual and co-cultured growth at the 5#x0025; level (paired *t-*test).

In the reciprocal experiment (1:100 *Psa* NZ54: *P. syringae* G33C), *Psa* NZ54 successfully invaded from rare in both the endophytic and epiphytic environment, although the rate of invasion was reduced on the leaf surface. However, both epi- and endophytic population sizes were significantly reduced compared to single inoculations (*P* <0.01, paired *t*-tests) (Figure 7). Despite the growing population of *Psa* NZ54, *P. syringae* G33C maintained the same epiphytic population size as when inoculated individually (*P* >0.2, paired *t*-tests). The endophytic population size of *P. syringae* G33C increased (*P* <0.01, 7dpi, paired *t*-test), which mirrored the results from the 1:1 competition experiments, where the presence of *Psa* NZ54 also had a positive effect on growth of *P. syringae* G33C.

In order to establish whether there was an advantage to being an early colonist, a time-stagger experiment was performed to see whether immigration history influences the interaction (Fukami *et al*., 2007). ‘Hort16A’ plants were pre-inoculated with either of the two strains and followed by a subsequent inoculation of the other strain after three days. Early colonization provided no advantage to *Psa* NZ54 (Figure 8A), as *P. syringae* G33C maintained and established a stable population by 7 dpi and exhibited no significant difference in growth compared to individual growth (*P* >0.3, paired *t*-tests). Reducing the secondary inoculation density resulted in a reduced initial population size at 3 and 7 dpi for *P. syringae* G33C (*P* <0.05, paired *t*-tests), but by 10 dpi the level was the same as when the two strains were grown individually (Figure S7C).

**Figure 8.**
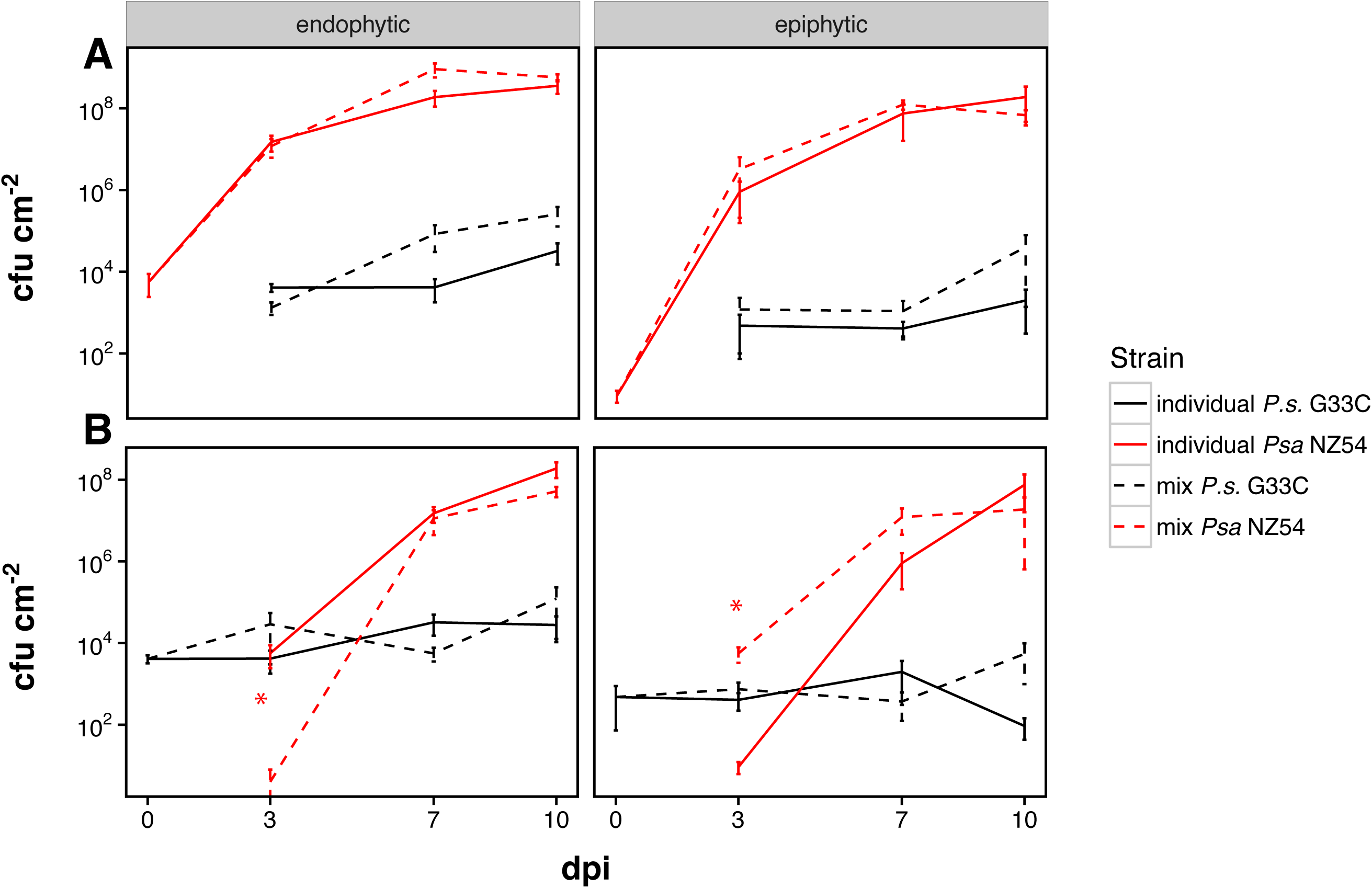
In planta priority effect of *Psa* NZ54 or P. syingae G33C with subsequent inoculation of the respective second strain with the same founding density. (A) *In planta* growth assay of *P. syringae* G33C using ‘Hort16A’ plantlets pre-inoculated for three days with *Psa* NZ54 (8×10^7^ cfu ml^−1^). (B) *in planta* growth assay of *Psa* NZ54 using ‘Hort16A’ plantlets pre-inoculated for three days with *P. syringae* G33C (8×10^7^ cfu ml_-1_). Solid lines represent individual growth and dashed lines represent growth in competition. The presented mean and standard error were calculated from the mean of five individual measurements. Asterisks indicate significance between individual and co-cultured growth at the 5#x0025; level (paired *t-*test).

When *P. syringae* G33C was the first colonist (Figure 8B) *Psa* NZ54 grew to the same population size by 7 dpi as compared to individually inoculated plants (*P* >0.5, paired *t*-tests). The growth of *Psa* NZ54 was initially lower compared to individual growth when inoculated at a lower density (*P* <0.05, paired *t*-tests), but this difference was no longer evident for the epiphytic population by 10 dpi (Figure S7B).

## DISCUSSION

Studies of pathogen populations rarely take into consideration co-occurring commensal types and yet such types are likely to be important contributors to population structure and infection progress (Lindow and Brandl, 2003; Demba Diallo *et al*., 2012; Bartoli *et al*., 2015; Buonaurio *et al*., 2015; Rufián *et al*., 2017). Here, with focus on P. syringae, we have combined traditional population genetic approaches with experiments designed to investigate interactions among members of an ecologically cohesive population. The most significant findings include (i) a clonal population structure for commensal kiwifruit *P. syringae* (ii) strong association of genetic diversity with ecological factors, (iii) discovery of a new clade of kiwifruit-associated kiwifruit *P. syringae* within PG3 (PG3a) (Figure 2, Figure 3), (iv) complex interactions between the pathogenic *Psa* isolate and PG3a with evidence of a stable polymorphism under some *in vitro* conditions, but not in planta (Figure 4, Figure 6).

Overall, we found that *P. syringae* from kiwifruit display a clonal population structure, comprised of two clonal complexes and a small number of abundant STs. This is in accordance with earlier reports of clonal population structure for P. syringae, despite focus on pathogenic isolates which tend to undergo clonal expansion upon host specialisation (Sarkar and Guttman, 2004). Homologous recombination events are few and limited to within phylogroups for P. syringae, which is also supported by well-defined phylogenetic clades (Baltrus *et al*., 2011; Bull *et al*., 2011; Berge *et al*., 2014; Nowell *et al*., 2016). A more fine-scale analysis of a collection of *Pseudomonas viridiflava* (now *P. syringae* PG7 and PG8 (Bartoli *et al*., 2014; Berge *et al*., 2014)) isolated from Arabidopsis thaliana suggests that recombination at the phylogroup level is primarily within-clade rather than between clade (Goss *et al*., 2005). Apart from the occurrence of recombination at the local scale, evidence of recombination has also been shown between crop strains and environmental isolates (Monteil *et al*., 2013).

Genetic diversity varied according to ecological factors, most strikingly for *P. syringae* collected from infected orchards, where genetic diversity was highest. This may reflect effects of *Psa* on the kiwifruit immune response, which may facilitate migration of leaf colonists into the apoplast and vascular tissues and thus allow access to water and nutrients. Such effects have been reported for infection of potatoes by *Pectobacterium atrosepticum* (Kõiv *et al*., 2015) and herbivore-damaged bitter cress leaves (*Cardamine cordifolia*) (Humphrey *et al*., 2014), where in both instances Pseudomonas population densities and diversity increased following plant damage.

We observed differences in *P. syringae* genetic diversity that appear to be attributable to differences in plant genotypes. Host species and cultivar identity is known to significantly affect the composition of phyllosphere bacterial communities (Adams and Kloepper, 2002; Van Overbeek and Van Elsas, 2008; Whipps *et al*., 2008; Bodenhausen *et al*., 2014; Laforest-Lapointe *et al*., 2016; Wagner *et al*., 2016). Differences in phyllosphere *P. syringae* diversity may also be influenced by environmental factors (such as humidity, nutrient availability or UV radiation) and orchard management practices. Different fertilizer and spray regimes (copper, antibiotics, Actigard^TM^ and biological agents) are employed by growers to prevent or manage *Psa* infection throughout the growing season (http://www.kvh.org.nz/vdb/document/99346). These practices may have selected for copper and streptomycin resistance in *Psa* and kiwifruit epiphytes in NZ and elsewhere (Han *et al*., 2003; Colombi *et al*., 2017; Petriccione *et al*., 2017).

Strains grouping with four major phylogroups (PG1, PG2, PG3, PG5) were recovered. This level of diversity in a cultivated environment is not surprising (Goss *et al*., 2005; Bull *et al*., 2011; Kniskern *et al*., 2011; Beiki *et al*., 2016; Hall *et al*., 2016). Two clades of endophytic *P. syringae* pv. syringae were recovered from symptomatic grapevines in Australia with pathogenic and non-pathogenic isolates clustering together (Hall *et al*., 2016). Samples obtained from citrus orchards suffering from citrus blast caused by *P. syringae* pv. syringae revealed isolates associated with PG2, PG7 and an unknown clade (Beiki *et al*., 2016). Two distinct and highly divergent subclades of Pseudomonas viridiflava (*P. syringae* PG7) were recovered from a global sampling of wild A. thaliana (Goss *et al*., 2005).

The newly recognised PG3a subclade of *P. syringae* appear to colonise kiwifruit leaves not only in NZ (dating back to 2010 (Visnovsky *et al*., 2016)), but also in other kiwifruit growing regions of the world, including Japan and China. Data from leaf samples indicate that PG3a is not displaced by *Psa*, but the total number of PG3a isolates collected is reduced in infected orchards. Interestingly the diversity of PG3a does not seem to be affected by infection status. Strains clustering with PG3a formed the majority (>50%) of kiwifruit isolates, and these have not yet been isolated from any other plant hosts recorded in PAMDB, with the exception of isolate 47L9, collected from tea leaves (*Camellia* sp.) growing in a former kiwifruit orchard. This indicates that PG3a forms a persistent association with kiwifruit plants – an observation that is further supported by the repeated isolation of PG3a from kiwifruit across large geographic distances, suggests that PG3a may have been coevolving with its host for some time. PG3a is thus also likely to be disseminated with the exchange of plant material (such as pollen or plant cuttings) between kiwifruit growing countries. The prevalence of PG3a in other kiwifruit growing countries (e.g. Korea, France or Italy) is at present unknown. The preferential occurrence of PG3 with woody hosts (Bartoli *et al*., 2015; Nowell *et al*., 2016) could explain the particular grouping of the kiwifruit resident clade within PG3. A similarly intriguing signal of host association was found in a collection of *P. syringae* isolates from A. thaliana, where PG2 representatives dominated (Kniskern *et al*., 2011; Karasov *et al*., 2017). Distinct lineages of non-pathogenic isolates have also been described for other plant pathogens such as *Xanthomonas arboricola*, where non-pathogenic strains are distant relatives of pathogenic lineages, despite being isolated from the same host (Essakhi *et al*., 2015; Triplett *et al*., 2015).

The kiwifruit commensal *P. syringae* G33C (representative of the PG3a subclade) successfully colonized the leaf surface and apoplast of kiwifruit without production of visible disease symptoms. This is reflected in the population size, which was reduced by 4-logs compared to pathogenic population size of *Psa* at 3 dpi. Similar population sizes have been reported for *P. syringae* pv. *phaseolicola*, which grows to a four-log higher population size on its host plant *Phaseolus vulgaris* compared to a non-pathogenic isolate; a 4-log reduction was also observed in resistant vs susceptible hosts (Omer and Wood, 1969; Young, 1974). Similar observations have been made for other plant pathogens, for example non-pathogenic *Xanthomonas* sp. displays a 4-log reduced growth compared to the disease-causing X. *oryzae* pv. *oryzae* (Triplett *et al*., 2015). Bacterial population density appears to be directly related to the production of disease symptoms, as was demonstrated for environmental *P. syringae* strains inoculated in kiwifruit, which grew to near pathogen population size levels, but induced symptoms of disease (Bartoli *et al*., 2015).

When grown in competition with *Psa* NZ54, *P. syringae* G33C population size increased. Studies exploring the dynamics of mixed infections have demonstrated inoculum density-dependent effects on colonization (Young, 1974; Macho *et al*., 2007). The ability of non-pathogenic *P. syringae* to colonize wider territories in the presence of a pathogenic strain was nicely demonstrated in a confocal microscopy study (Rufián *et al*., 2017). It is possible that *P. syringae* G33C benefits from the virulence activity of *Psa* NZ54. T3SS-dependent hitch-hiking effects have been observed for *P. syringae* pv. *syringae* (Hirano *et al*., 1999), however the increase in *P. syringae* G33C growth persists even in the absence of a functional T3SS in the pathogenic *Psa* strain. Virulence activities not encoded by the T3SS, such as phytotoxin production, may be responsible for this outcome.

Epiphytic and *in vitro* growth of *Psa* NZ54 was significantly reduced when co-inoculated with *P. syringae* G33C. Similarly, *Psa* growth may be suppressed by co-inoculation with environmental isolates of *P. syringae* (Bartoli *et al*., 2015). Epiphytes may suppress pathogen growth either as direct antagonists or indirectly via resource competition (Wilson and Lindow, 1994). The specific mechanism by which *P. syringae* G33C suppresses *Psa* remains undetermined. The *Psa* NZ54 population collapse in shaken M9 was delayed at 10:1 inoculation ratios, which suggests that this effect was most likely due to the accumulation of antimicrobial compounds produced by *P. syringae* G33C. Phytotoxin production is widespread among fluorescent pseudomonads with some toxins having antimicrobial activity that can be induced dependent on culture conditions (still vs. shaken) *in vitro* (Durbin, 1982; Bender *et al*., 1999). Contact-dependant growth inhibition (CDI) via Type 5 and 6 secretion systems or bacteriocins may also mediate *P. syringae* interactions (Hayes *et al*., 2010; Haapalainen *et al*., 2012; Ruhe *et al*., 2013; Hockett *et al*., 2015).

Our in-depth localised sampling has revealed a global association of PG3a with kiwifruit. Additionally, we have shown that this clade of non-pathogenic *P. syringae* engage in complex interactions with pathogenic *Psa*. This highlights the value of understanding genotypic diversity and ecological interactions among pathogens and non-pathogens in field settings. Clarifying how commensals persist in association with specific hosts over long periods without causing disease and the mechanism by which they modulate pathogen invasion and proliferation will contribute to a fuller understanding of plant-microbe interactions.

## EXPERIMENTAL PROCEDURES

### Plant tissue collection and bacterial isolation

The sampling scheme was designed to obtain strains from *Psa* infected and uninfected hosts, irrespective of disease stage (symptom development) or pathogenicity potential of the isolate. *P. syringae* was isolated from the leaf surfaces of two different cultivars of *Actinidia chinensis*: *A. chinensis* var. *chinensis* Hort16A (gold) and A. *chinensis* var. *deliciosa* Hayward (green), which vary in their susceptibility to *Psa*: ‘Hort16A’ is more susceptible than the green ‘Hayward’ (Ferrante and Scortichini, 2010; Cameron and Sarojini, 2014). One infected and one uninfected orchard of each variety was sampled by collecting three leaves from six separate vines along a diagonal path of ∼400m (Table S1). Sampling occurred at three intervals during the growing season: spring (after bud break), summer and autumn (prior to harvest). Vine trunks and canes (secondary branches) were tagged to ensure resampling of the same spot. Some ‘Hort16A’ canes were removed during routine disease management; neighbouring canes on the same vine were then sampled and tagged. All uninfected Hayward vines were cut down prior to the last sampling day so the adjoining block of Hayward was sampled instead. The location of each sampled orchard is listed in Table S2.

Leaves were individually placed in 50 mL conical centrifuge tubes and washed with 40 mL 10mM MgSO_4_ buffer supplemented with 0.2 #x0025; Tween (Invitrogen, US) by alternately shaking and vortexing at slow speed for 3 min. After removing the leaf, the leaf wash was centrifuged at 4600 rpm for 10 min. The supernatant was removed and the pellet was resuspended in 200 μL 10mM MgSO_4_. 100 μL of the resuspension was stored at −80°C and 100 μL was plated on *Pseudomonas* agar base (Oxoid, UK) supplemented with 10 mg/L cetrimide, 10 mg/L fucidin and 50 mg/L cephalosporin (CFC supplement, Oxoid, UK). For each leaf two isolates exhibiting *P. syringae* colony morphology (round, creamy white) were selected randomly from the plate and restreaked, then used to inoculate liquid overnight cultures for storage at −80°C. Isolates were tested for the absence of cytochrome C oxidase using Bactident Oxidase strips (Merck KgaA, Germany), characteristic of P. syringae. A total of 148 *P. syringae* isolates were obtained from the four orchards (Table S1).

### PCR amplification & sequencing

A lysate was prepared for each isolate by resuspending a colony in 100 μL ddH_2_0 and lysing the cells at 96°C for 10 min. Strains were sequenced using the Hwang *et al*. (2005) MLST scheme for four housekeeping genes *gapA, gyrB, gltA (=cts)* and *rpoD* (reverse). Due to amplification problems, the forward primer for *rpoD* from Sarkar and Guttman (2004) was used. PCR amplification was performed with a BIO-RAD T100 Thermal Cycler following an adapted protocol of Hwang *et al*. (2005): a total reaction volume of 50 μl with a final concentration of 1x PCR buffer (Invitrogen, US), 1 μM for each primer, 0.2 mM dNTP’s (Bioline, UK), 1 U Taq Polymerase (Invitrogen, US), 1 μl lysed bacterial cells, 2% DMSO (Sigma-Aldrich, US) and 1.5 mM MgCl_2_. Initial denaturation was at 94°C for 2 min, followed by 30 cycles of amplification with denaturation at 94°C for 30 s, annealing at 63°C for 30 s and elongation at 72°C for 1 min. Final elongation was for 3 min at 72°C. Samples were purified using the Exo-CIP method and sequenced by Macrogen Inc (South Korea). Sequence analysis was performed with Geneious v7.1.7 (Kearse *et al.*, 2012). Sequences were trimmed to the same length (476 bp *gap1*, 507 bp *gyrB*, 529 bp *gltA*, 498 bp *rpoD*) and concatenated (201 0 bp) (GenBank accession numbers: *gapA* MG642149 - MG642296; *gyrB* MG642297 - MG642444; *gltA* MG642445 - MG642592; *rpoD* MG642593 - MG642740).

### Population genetics

#### Sequence diversity indices

A rarefaction analysis was performed using MOTHUR v.1.34.4 by subsampling using 1,000 iterations (Schloss *et al.*, 2009). Pairwise genetic distances between isolates were calculated and sequences assigned to Operational Taxonomic Units (OTUs) based on the corresponding average pairwise genetic distance of each group.

Simpson’s index of diversity (D) and evenness (ED) (α-diversity) and Sørensen’s index of dissimilarity (β-diversity) were calculated using the *vegan* package (Oksanen *et al.*, 2016) in R v3.3.1 (R.Core.Team, 2016). Simpson’s D was converted to the effective number of species (D_c_) in order to account for the non-linear properties of Simpson’s index of diversity (Jost, 2006).

#### Multilocus Sequence Typing

Sequence types (STs) sharing three out of four alleles (SLV, single locus variants) were grouped using eBURST v3 (bootstrapped with 1,000,000 resamplings) (Feil *et al.*, 2004; Spratt *et al.*, 2004). A Minimum Spanning Tree providing an overview of triple locus variants was constructed using Phyloviz v2.0 (Francisco *et al.*, 2012).

165 non-redundant ST profiles of *P. syringae* strains were downloaded from the Plant Associated and Environmental Microbes Database (PAMDB, http://genome.ppws.vt.edu/cgi-bin/MLST/home.pl) (Almeida *et al.*, 2010). *P. syringae* sequences isolated recently from kiwifruit and air in Japan (NCBI) and kiwifruit isolates from NZ, France and the United States (Visnovsky *et al.*, 2016) were also included. A reduced set of 37 *P. syringae* isolates representing the different monophyletic groups of *P. syringae*, as well as the Japanese kiwifruit strains and the US, France and NZ kiwifruit isolates from previous years were used to provide better resolution in the phylogenies displayed in Figure 2, Figure 3 and Figure S4.

#### Sequence diversity and recombination

START2 (v0.9.0 beta) was employed to calculate parameters of genetic diversity, number of alleles and polymorphic sites, GC content and the ratio of non-synonymous to synonymous substitutions (*d*_*N*_/*d*_*S*_ ratio) (Jolley *et al.*, 2001). The number of mutations and amino acid changes and nucleotide diversity parameter π were calculated with DnaSP v. 5.10.1 (Rozas and Rozas, 1995). Jmodeltest 2.1.7 (Guindon and Gascuel, 2003; Darriba *et al.*, 2012) was used with default parameter settings to find the best-fitting evolutionary model. Pairwise genetic variability among and between phylogroups was calculated using MEGA7 (Kumar *et al.*, 2016).

To test whether genetic diversity varied by sampling location, time of sampling, orchard infection status and/or cultivar, a permutational multivariate analysis of variance (PERMANOVA) (Anderson, 2001; McArdle and Anderson, 2001) was performed using PRIMER v 6.1.12 (PRIMER-E Ltd., Plymouth, UK, PERMANOVA+ add-on v. 1.0.2.). Pairwise distances among unique STs were used as input and tests were run with 9999 permutations. Any effects due to unequal sample size were taken into account using the Type III SS (sum of squares) option.

LDHAT v2.2a (Auton and McVean, 2007) was used to estimate the rate of mutation (Watterson’s θ) and recombination (ρ) using the composite likelihood method of Hudson (Hudson, 2001) with an adaption to finite-site models. Only polymorphic sites with two alleles were included and the frequency cut-off for missing data was set to 0.2.

#### Phylogenetic reconstruction

Trees were built using single representatives of each unique ST from this study to improve readability of the tree. MrBayes v.3.2 (Ronquist *et al.*, 2012) was used to construct Bayesian trees, using the best-fitting evolutionary model (jModeltest) for individual genes and the concatenated alignment. Single gene trees (Figure S4) were constructed with TREEPUZZLE (Schmidt *et al.*, 2002) and Dnaml (PHYLIP v3.695, Felsenstein 1989) was used to test for congruence between single trees (SH-test) using default parameters, providing Maximum Likelihood trees as input and a random number seed of 333.

### Strains and culture conditions

A list of all bacterial strains used in this study can be found in Table S2. *Pseudomonas* strains were cultured in King’s B or minimal M9 media at 28°C and *E. coli* was cultured in Luria Bertani medium at 37°C. Liquid overnight cultures were inoculated from single colonies and shaken at 250 rpm for 16 hrs. The antibiotics kanamycin (kan) and nitrofurantoin (nf) were used at a concentration of 50 µg/ml. Kanamycin resistant *Psa* NZ54 and *Psa* NZ13 Δ*hrcC* were employed in all *in vitro* and *in planta* experiments. Both *Psa* strains belong to the same pandemic clade of *Psa* (biovar 3), and differ in three SNPs across the core genome as defined in McCann *et al.* (2017).

### Mutant development

*Psa* NZ13Δ*hrcC* was constructed by in-frame deletion of *hrcC* via marker exchange mutagenesis. Knockout construct was generated by overlap extension PCR (Ho *et al.*, 1989) using the primers listed in Table S6. DNA was amplified from *Psa* NZ13 with Phusion® High-Fidelity DNA polymerase. The deletion construct was inserted into pK18mobsacB (Schäfer *et al.*, 1994). The recombinant vector was transferred into *Psa* NZ13 via triparental mating, using as helper *E. coli* DH5α strain containing pRK2013. Mutants were selected by plating on KB kanamycin (50 µg/mL) and subsequently on KB containing 5#x0025; sucrose. Mutants were screened by PCR using external primers (Table S6) and the deletion was then confirmed by sequencing.

Triparental matings were performed to introduce a kanamycin resistant Tn*5* transposon into *Psa* NZ54 and *Psa* NZ13 Δ*hrcC*. *E. coli* S17-1 Tn*5hah Sgid1* (donor) (Zhang *et al.*, 2015), *E. coli pRK2013* (helper) (Ditta *et al.*, 1980) and *Psa* NZ54 or *Psa* NZ13 Δ*hrcC* (recipient) were grown in shaken liquid media overnight. 200 µ l of donor and helper and 2mL of recipient were individually washed, pelleted and combined in 30 µ l 10 mM MgCl_2._ The mixture was plated on a pre-warmed LB agar plate and incubated at 28°C for 24 hrs. The cells were scraped off and resuspended in 1 mL 10 mM MgCl_2_ and plated on KB plates supplemented with kanamycin and nitrofurantoin. Bacterial growth was compared to the wild type recipient in both KB and M9 media to ensure marker introduction did not result in a loss of fitness.

### Competition assays

The two isolates (*P. syringae* G33C and *Psa* NZ54) used for competition assays were isolated from the same leaf in a *Psa* infected orchard, reflecting co-occurrence of the strains in nature.

#### In vitro competition assays

Competition experiments were performed *in vitro* using rich (King’s B) and minimal (M9) media in a shaken and static environment. Competition experiments were performed in 1:1, 1:10 and 10:1 ratios for each of the four assay conditions. Liquid overnight cultures of each strain in KB were established from single colony inoculations. 30 mL vials with 4 mL of the appropriate media were inoculated with each strain, adjusted to a founding density of either 5×10^6^ cfu ml^−1^ (OD_600_ 0.006) or 4×10^4^ cfu ml^−1^ (OD_600_ 0.0004). Control vials were inoculated with a single strain, adjusted to 5×10^6^ cfu ml^−1^. Cultures were incubated at 28°C and grown over a period of 72 hrs, either still or shaken at 250rpm. Bacterial density was calculated at 0, 24,48 and 72 hrs by plating dilutions on KB kan and M9 agar plates to distinguish between competing strains. The experiment was performed using three replicates and repeated three times.

#### In planta competition and pathogenicity assays

Epiphytic and endophytic growth of *Psa* NZ54, *Psa* NZ13 Δ*hrcC* and *P. syringae* G33C was evaluated on 4-week old kiwifruit plantlets using single and mixed-culture inoculation. Clonally propagated *A. chinensis* var. *chinensis* ‘Hort16A’ and ‘SunGold’ were grown for a minimum of one month in a Conviron CMP6010 growth cabinet at 21°C with a 14/10 hr light/dark cycle and 70#x0025; humidity. Bacterial strains were incubated for two days at 28°C on KB plates, after which they were resuspended in 10 mM MgSO_4_ buffer. Mixed inoculum (1:1, 1:100 and 100:1) was prepared in 50 mL 10 mM MgSO_4_ buffer and 0.002% Silwet-70 (surfactant), with strains adjusted to 8×10^7^ cfu ml^−1^ (OD_600_ 0.1) or 8×10^5^ cfu ml^−1^ (OD_600_ 0.001). Single strain plant inoculations were also performed using an initial 8×10^7^ cfu ml^−1^ (OD_600_0.1).

Plants were inoculated by submerging leaves in the inoculum for 5 s and allowing to air-dry. Plants were returned to the growth cabinet and watered every second day. Bacterial density was assessed at either 0, 2, 4, 7 and 10 days post inoculation (dpi) or 0, 3, and 7 dpi (Δ*hrcC* competition experiments). Epiphytic growth was assessed by placing inoculated leaves in separate sterile plastic bags with 35 mL 10 mM MgSO_4_ buffer and shaking manually for 3 minutes. The leaf wash was centrifuged at 4600 rpm for 3 min and the supernatant discarded. Bacteria were resuspended in 200 µl buffer and serial dilutions plated on M9 and KB+kan agar plates.

Endophytic growth was assessed by removing one 1cm^2^ leaf disk per plant (including the midrib), surface sterilizing in 70% EtOH for 30 sec, drying and homogenising for 1 minute in a 1.5 mL Eppendorf tube containing 200 μl buffer and two metal beads with the TissueLyser II (QIAGEN). The plant homogenate was serially diluted and plated on M9 and KB+kan agar plates. All experiments were performed in duplicate, with at least 4 replicates per experiment.

### Statistical analysis

A Student’s *t*-test was used to verify the statistical difference where applicable. For non-normally distributed data with unequal variance, the Mann Whitney U test was performed.

The fitness of each strain in the competition experiments is expressed as the Malthusian parameter (Lenski *et al.*, 1991). The Malthusian parameter was calculated as *M*= (*ln*(*N*1_*f*1_/*N*1_*i*_)) /(*ln*(*N*2_*f*_/*N*2_*i*_)), where *N*1_*i*_ is initial number of cfu of strain 1 at 0h and *N*1_*f*_ cfu after 24 hrs (*in vitro*) or 2/3 dpi (*in planta*, ‘Hort16A’/’SunGold’).

## BIOSECURITY AND APPROVAL

All worked was performed in approved facilities and in accord with APP201675, APP201730, APP202231.

## ACKNOWLEDGEMENTS

We want to particularly thank kiwifruit growers David French, Rex Reed and Bruce and Fiona Aitken for granting access to their orchards. The work conducted was supported by the New Zealand Ministry for Business, Innovation and Employment (C11X1205). Christina Straub was supported through a PhD scholarship from the New Zealand Institute for Advanced Studies (NZIAS) and Plant & Food Research.

## SUPPORTING INFORMATION

**Figure S1. Rarefaction curves based on the concatenated sequences.** Two curves each are shown for *P. syringae* (n=148) and the sequences grouped according to phylogroups (PG): solid lines represent grouping based on unique STs and dashed lines according to a cut-off equal to the average pairwise genetic distance of the group: PG1, PG2 & PG3 = 0.02 cut-off, Psyr all = 0.05 cut-off.

**Figure S2. Shared and unique STs among orchards.** Gold I = infected ‘Hort16A’; Gold NI= uninfected ‘Hort16A’; Green NI = uninfected ‘Hayward’; Green I = infected ‘Hayward’ orchard; n= number of STs found in orchard.

**Figure S3. eBurst snapshot of STs at the single locus variant level.** The size of the circles correlates with the frequency of the respective ST found in the dataset. Colours correspond to the different orchards, orange = infected ‘Hort16A’, yellow = uninfected ‘Hort16A, dark green = infected ‘Hayward’, light green = uninfected ‘Hayward’. CC = clonal complex.

**Figure S4. Maximum Likelihood trees based on single genes.** Each Maximum Likelihood tree is rooted on *Pseudomonas fluorescens* SBW 25 and was reconstructed using TREEPUZZLE based on the Tamura-Nei model using 100,000 puzzling steps. Trees were built using single representatives of each unique ST to improve readability of the tree. Values indicated at nodes are bootstrap values. The corresponding phylogroup distinctions based on the concatenated ML tree are indicated with the coloured squares.

**Figure S5. Leaves of ‘Hort16A’ plants inoculated with *Psa* NZ54, *P. syringae* G33C and a 1:1 mix of *Psa* NZ54: *P. syringae* G33C at 2, 4, and 7 days post inoculation.**

**Figure S6. Leaves of ‘SunGold’ plants inoculated with *Psa* NZ54, *P. syringae* G33C, *Psa* NZ13 ΔhrcC and 1:1 mix of the respective strain combinations at 3 and 7 days post inoculation**. For leaves showing minor leaf spots, the lower side of the leaf is also shown for easier detection of symptoms.

**Figure S7. *In planta* priority effect of *Psa* NZ54 or *P. syringae* G33C with subsequent inoculation of the second strain with 100-fold lower concentration.** *In planta* growth assay using ‘Hort16A’ plantlets pre-inoculated (8×10^7^ cfu ml^−1^) with one strain followed by inoculation of the second strain at (8×10^5^ cfu ml^−1^). The two panels display growth curves for endo- and epiphytic growth respectively. A) Individual growth, B) inoculation of *Psa* NZ 54 at day 3 and C) inoculation of *P. syringae* G33C at 3 dpi. Solid lines represent individual growth and dashed lines represent growth in competition. The presented mean and standard error were calculated from the mean of five individual measurements. Asterisks indicate significance between individual and co-cultured growth at the 5% level (paired *t-*test).

**Table S1. Geographic location of orchards, strain summaries and diversity indices per orchard.** Specification of cultivar and infection status at the time according to KVH (Kiwifruit Vine Health), orchard ID, GPS coordinates, location and month of sampling, N = number of collected *P. syringae* strains, N PG3a = number of PG3a strains in total sample, N STs = number of unique STs, N STs PG3a = number of unique STs grouping with PG3a, D = Simpsons index of diversity, D_c_ = converted to effective number of species, ED = Simpsons evenness.

**Table S2. List of all strains.** Strain information and assigned sequence type of strains used for MLST study (all) and strains used for phylogenetic analysis (highlighted in grey). Phylogroup association only provided for isolates used for phylogenetic analysis. Alias provides the name used for competition experiments.

**Table S3. PERMANOVA results of 3-factor nested analysis for differences in genetic diversity.**

**Table S4. LDhat recombination analysis for host and disease status.** Length of alignment in bp, number of sequences, number of segregating sites, mutation rate θ, recombination rate ρ and ratio ε (ρ/ θ).

**Table S5. LDhat recombination analysis for global data sorted according to phylogroup (PG)**. Length of alignment in bp, N = number of sequences, number of segregating sites, mutation rate θ, recombination rate ρ and ratio ε (ρ/ θ).

**Table S6. List of primers used for construction of the deletion mutant *Psa* NZ13 Δ*hrcC*.**

